# EBNA1 INHIBITORS REVEAL CDC7 AND POU2F1 AS DIRECT FUNCTIONAL TARGETS IN EBV EPITHELIAL CANCERS

**DOI:** 10.64898/2026.02.05.703959

**Authors:** Songtao He, Niseno Terhuja, Samantha Soldan, Christopher Chen, Joel Cassel, Xiangfan Yin, Qin Liu, Sun Sook Chung, Leonardo Josué Castro-Muñoz, Leena Yoon, Jie Wang, Joseph M. Salvino, Benjamin Gewurz, Italo Tempera, Troy E. Messick, Paul M. Lieberman

**Author notes:** Corresponding Author Paul M. Lieberman, The Wistar Institute, Philadelphia, PA 19104.

## Abstract

Epstein-Barr virus (EBV) latent infection is causally linked to several epithelial cancers, including endemic forms of undifferentiated nasopharyngeal carcinoma (NPC) and to a subtype of gastric cancer (GC). EBNA1 is the viral-encoded sequence-specific DNA-binding protein required for episome maintenance but also contributes to host-cell survival through multiple mechanisms including binding to host chromosome. We previously developed small molecule inhibitors of EBNA1 DNA-binding that block host cell cycle progression and growth of EBV+ tumors *in vivo*. However, the underlying molecular mechanisms of EBNA1 function and inhibition have not been completely elucidated. In this study, we employ VK1727 to inhibit EBNA1 DNA-binding to viral and cellular genomes in three EBV+ epithelial tumors (PDX C15, C666-1 and SNU719). We integrate EBNA1ChIP-seq and transcriptomic RNA-seq analyses to identify the cell cycle dependent kinase CDC7 and a stem cell transcription factor POU2F1 as direct functional targets of EBNA1 in these epithelial cancers. EBNA1 binding to CDC7 promoter and POU2F1 intron promotes RNA Pol II-pS5 to initiate transcription of these two genes. We show that CDC7 inhibitor Simurosertib is epistatic, while Bcl2 inhibitor Venetoclax is synergistic with VK1727 in the inhibition of EBV+ epithelial cancer cell proliferation and survival. Our study reveals new functional gene targets and pathways of VK1727 in EBV+ epithelial cancers that provide new biomarkers and combinatorial strategies to treat EBV-driven cancers.

**IMPORTANCE:** EBNA1 is essential for EBV latency and tumorigenesis, but its mechanism of action on host gene expression is not yet known. Small molecule inhibitors of EBNA1 DNA-binding block cell cycle progression and inhibit growth of EBV+ tumors. In this study, we use the EBNA1 small molecule inhibitor VK1727 to identify cellular gene targets that are bound by EBNA1 and deregulated by its pharmacological inhibition in EBV+ epithelial cancer cell lines and an NPC PDX mouse model. We identify cycle dependent kinase CDC7 and the stem cell transcription factor POU2F1 as EBNA1 bound and regulated genes important for EBV epithelial cancer proliferation. These findings not only decipher molecular mechanism how VK1727 blocks cell cycle progression and inhibits cell proliferation but also provide two new cellular gene targets and pathways for therapeutic intervention in EBV+ epithelial cancers.

## INTRODUCTION

Epstein-Barr virus (EBV) is a human gammaherpesvirus that establishes lifelong latent infection in more than 95% of the human population (1). EBV is also classified as a tumor virus due its strong association with several human cancers, including Burkitt’s lymphoma (BL), Hodgkin’s disease (HD), gastric (GC) and nasopharyngeal carcinoma (NPC) (2). EBV epithelial cancers represent ∼70% of all EBV-associated cancers worldwide (3). NPC is highly prevalent in southern China, Southeast Asia and some North African countries with an incidence of 4–25 cases per 100,000 individuals (4). EBV-associated gastric cancer (EBVaGC) is a specific molecular subtype comprising ∼10% of all gastric cancers world-wide (5). EBV DNA and viral gene products are readily detected in EBV epithelial cancers, but the patterns of gene expression and the targets of oncogenic transformation in epithelial tumors are generally different than in EBV-associated lymphomas. While EBV can efficiently immortalize B-lymphocytes in cell culture and cause B-cell lymphomas in immunosuppressed individuals (6, 7), its mechanism of oncogenic transformation in epithelial cancers is less well understood.

EBV infection of normal epithelial cells results in a transient and lytic replication cycle (8). However, in EBV epithelial cancers, EBV maintains a latent state that retains episomal DNA in tumor cells (9, 10). Most epithelial cancers express a type II latency, characterized by the limited expression of only EBNA1 and LMP1/2 proteins, and many non-coding RNAs including EBERs and BARTs (11). LMP1 is a well characterized viral oncogene that functions as a viral mimic of the TNFR family member, CD40, that activates signaling pathways related to cellular growth and survival (12). LMP2 mimics BCR signaling in B-lymphocytes, and related Src-family kinase pathways in epithelial cells (13, 14). The EBV non-coding RNAs, including many miRNAs are known to have diverse activities that could contribute to viral oncogenesis (15). EBNA1 is ubiquitously expressed in most EBV-positive tumor cells in NPC and GC (16). Numerous studies have described the versatility of EBNA1 in maintaining episome structure, promoting viral replication, tethering episome to metaphase chromosomes, which is essential for maintaining EBV latency, regulating host gene expression and interacting with host proteins involved in viral sensing and cancer signaling pathways (16–18). Increasing evidence reveals an oncogenic role of EBNA1 in EBV associated epithelial cancers, such as promoting EMT (Epithelial-Mesenchymal Transition) process (19), inducing genomic instability (20), and enhancing immune evasion (21). While EBNA1 has been implicated in each of these oncogenic pathways, the specific molecular mechanisms require further investigation.

In addition to its direct binding to the EBV episome, EBNA1 can also bind to many sites in the host genome (22, 23). Previous studies from our lab characterized some EBNA1-bound gene targets in cellular genome, such as MEF2B, EBF1, IL6R in BL cell lines (24), and gastrokines GKN1 and GKN2 in EBVaGC cell lines (25). Others have found EBNA1 regulates NOX2 in EBVaGC cell lines to modulate ROS (26), and induce telomere dysfunction (27), and BMP2 in NPC (28). EBNA1 can also activate viral genes, such as EBNA2 in type III latency (29), and this is thought to be mediated by long-distance enhancer-like DNA-loop interactions and other epigenetic effects (30–32). However, the regulation of cellular gene expression by EBNA1 remains subtle and complex compared to the more potent transcriptional activation by EBNA2 and EBNA3C (33, 34).

EBNA1 has been considered an attractive target for small molecule inhibition due to its essential role in maintaining EBV episomes during latency, its consistent expression in cancer cells, and its unique and druggable protein structure (35, 36). A series of small molecule inhibitors have been developed using a structure-based drug design approach and shown to bind directly to EBNA1 and sterically inhibit DNA binding (36). VK1727 has been shown to reduce EBNA1 DNA-binding to oriP DNA *in vivo* and reduce viral copy number in EBV associated tumors (37). A very closely related analogue to VK1727 (e.g. VK2019) has been investigated in a clinical trial to treat NPC (38). Both VK1727 and VK2019 reduce EBV persistence in epithelial cancer cells and prevent tumor growth in mouse models of NPC and EBVaGC *in vivo* (37, 39). In addition, our lab demonstrated VK1727 specifically blocks cell cycle progression of EBV associated tumors (37). In this work, we investigate the mechanism through which the EBNA1 inhibitor VK1727 blocks cell cycle progression of EBV associated epithelial cancer cells. We assayed the effect of VK1727 on the genome-wide DNA binding of EBNA1 by ChIP-seq, and the transcriptional response by RNA-seq in EBVaGC line SNU719, and EBV+ NPC cell line C666-1, and in the C15 PDX mouse model of NPC. We integrated these data sets to identify common gene targets of VK1727. We characterized two functional gene targets of EBNA1 (CDC7 and POU2F1) in EBV+ epithelial cancers, which involve in cell cycle progression and cell fate determination. Our results here provide two potential biomarkers for evaluating the therapeutic activity of EBNA1 inhibitors in EBV+ epithelial cancers and new pathways for synergistic drug combinations.

## RESULTS

### EBNA1 inhibitors target cell cycle control in three different EBV tumor models

Our lab previously reported EBNA1 inhibitor VK1727 selectively blocks cell cycle progression and tumor growth of EBV+ epithelial cancers (37). But the underlying molecular mechanism remains unclear. To investigate further, we treated three EBV tumor models with VK1727 at short time (PDX C15, 5 days; C666-1, 2 days; SNU719, 2 days) and performed RNA-seq analysis. Transcriptomic analysis identified differentially expressed genes after VK1727 treatments for each cell line or PDX (Fig.1A-D). For PDX C15, there were 763 differentially expressed genes after VK1727 treatment (Fig. 1A). The top four enriched signaling pathways included cell cycle, TNF signaling pathway, cellular senescence, and P53 signaling pathway (*P* value <0.05, Fig.S1 A). Cell cycle was ranked as top one signaling pathway, with an enrichment score 0.6 (Fig.S1B), which suggested more upregulated genes than the downregulated genes being enriched in cell cycle (Fig.S1C). Transcriptomic and function analysis of VK1727 treated C666-1 cells identified 865 differentially expressed genes, primarily associated with P53 signaling pathway, cell cycle regulation, cellular senescence and TNF signaling (*P* value < 0.05, Fig.1B and Fig.S2A). The cell cycle pathway was identified as the second most significantly altered signaling pathway with a score -0.4 (Fig.S2B). This negative score indicates that a greater proportion of genes in the cell cycle pathway were downregulated rather than upregulated following VK1727 treatment (FigS2C). RNA-seq analysis of VK1727 treated SNU719 cells revealed 3249 differentially expressed genes, enriched in lipid metabolism and atherosclerosis pathways, cell cycle regulation, efferocytosis, AMPK signaling pathway, and P53 signaling pathway (*P* value < 0.05, Fig.1C, Fig.S3A). Cell cycle again was listed as the second most significantly altered signaling pathway with a score 0.4 (Fig.S3B and C). After that, we performed overlap analysis of differentially expressed genes among three VK1727 treated tumor models. 27 differentially expressed genes were enriched among three tumor models (Fig.1D). Function analysis demonstrated that these 27 overlap genes were enriched for cell cycle, cellular senescence, transcriptional dysregulation in cancer, microRNAs in cancer and P53 signaling pathway (*P* value < 0.05, Fig1E and Table S1).

**Figure 1.**
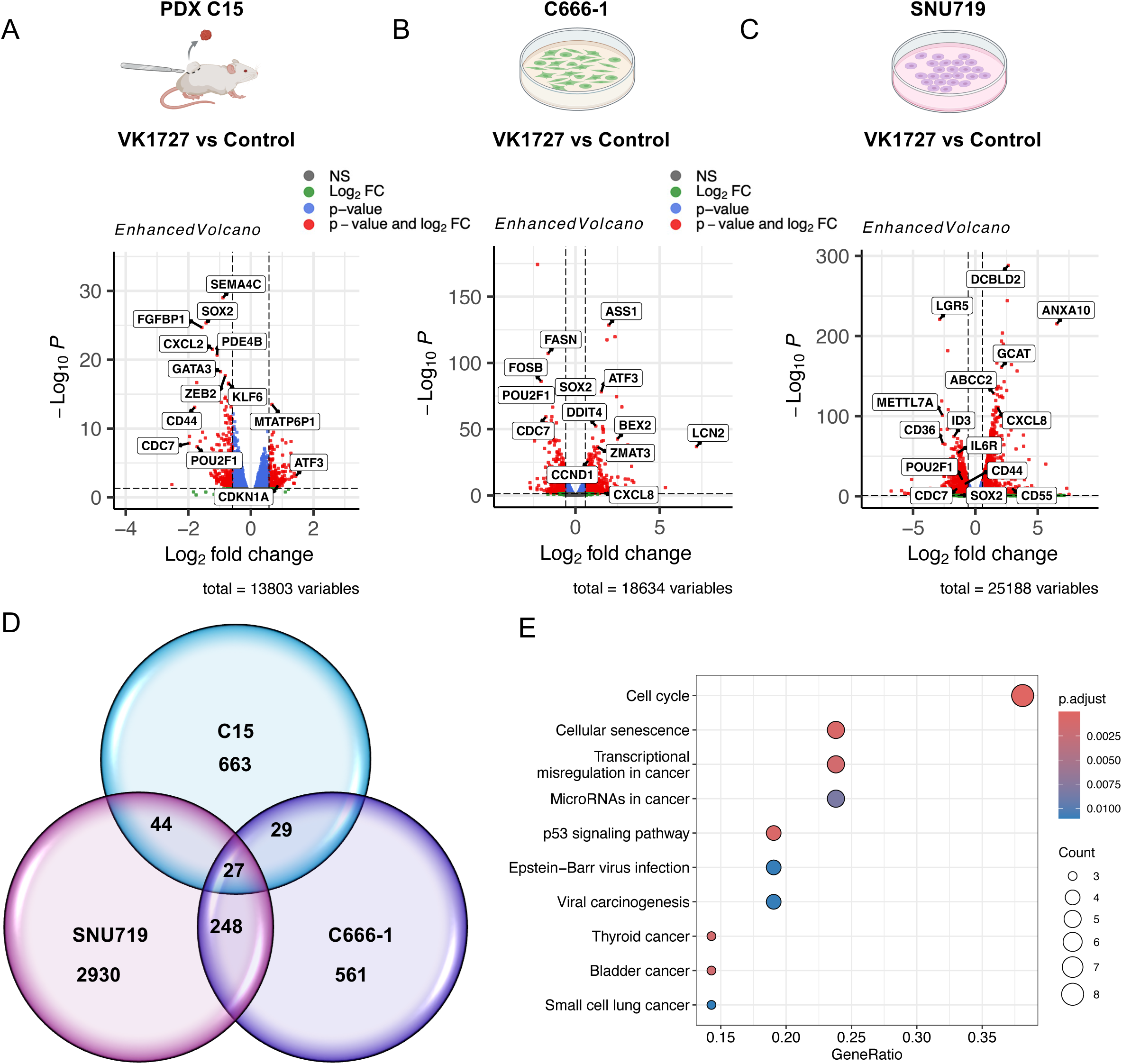
RNAseq analysis of VK1727 in three different EBV tumor models. **A)** Volcano plot of differentially expressed cellular genes in VK1727 treated PDX C15. 763 differentially expressed genes were identified with statistical significance (fold change >0.58 and p value <= 0.05). Red dots represent differentially expressed genes with fold change >0.58 and p value <= 0.05, green dots represent fold change >0.58 but p value >0.05, blue dots represent fold change <0.58 but p value <0.05 and black dots are genes that do not have significant expression change, with fold change <0.58 and p value >0.05. X-axis numerical scale labels are 2-base logarithm of fold change. Y-axis numerical scale labels are minus form of 10-base logarithm of P-value. **B)** Volcano plot of differentially expressed cellular genes in VK1727 treated C666-1 cells. RNA-seq analysis characterized 865 differentially expressed genes with statistical significance (fold change >0.58 and p value <= 0.05). **C)** Volcano plot of differentially expressed cellular genes in VK1727 treated SNU719 cells. RNA-seq analysis of VK1727 SNU719 cells identified 3249 differentially expressed genes with statistical significance (fold change >0.58 and p value <= 0.05). **D)** Venn diagram of overlap genes among three VK1727 treated tumor models. Differentially expressed genes were enriched with statistical significance (fold change >0.58 and p value <= 0.05) in each VK1727 treatment. The *match* function in R programing was used to enrich overlap of downregulated and upregulated genes among three tumor models. **E)** Function analysis of overlap genes among three VK1727 treated tumor models. The clusterProfiler package was employed to annotate 27 overlap genes into KEGG (Kyoto Encyclopedia of Genes and Genomes) database. DOSE and ggplot2 were used to generate dot plot of enriched signaling pathways. Count denotes the number of enriched genes in each signaling pathway. GeneRatio is the ratio of Count to setSize (total number of reported genes in each pathway). Adjusted *P* value was employed to assess statistical significance of an enriched signaling pathway.

We also performed differential expression analysis of EBV genes across the three tumor models. In VK1727 treated PDX C15 tumors, only a few viral genes were affected and the observed fold change was modest (< 1.5) (Fig.S4A). These include up-regulation of non-coding RNA EBER2 and RPMS1, and down regulation of EBNA1, LMP1, BNRF1, and BPLF1. VK1727 treated C666-1 also showed modest fold changes (< 1.5) for 11 genes, including upregulation of EBER2, RPMS1, BHLF1 and down regulation of EBNA1, LMP1, and EBNA3A (Fig. S4B). VK1727 treated SNU719 cells affected 22 EBV genes including upregulation of EBER1 and 2, RPMS1, LF1, LF2, LF3, and down regulation of EBNA1, LMP2A, BRLF1, BMRF1, and BPLF1 (Fig. S4C). Collectively, these findings indicate that VK1727 perturbs EBV gene expression but to different extents and patterns in different cell types.

### Integration of EBNA1 ChIP-seq with VK1727 RNA-seq to identify direct targets of EBNA1

To investigate direct targets of VK1727 in 27 differentially expressed genes, we re-analyzed previously published EBNA1 ChIP-seq datasets in MutuI, SNU719 and C666-1 cells (23, 24, 30). ChIP-seq analysis identified conservative EBNA1 binding sites in FR, DS and Qp regions in EBV genomes in three cell lines (Fig.2B), which confirmed reliability of ChIP-seq analysis. We next performed overlap analysis of EBNA1 ChIP-seq datasets between C666-1 and SNU719 cells. Using bedtools, we identified 318 conservative binding sites in cellular genomes in C666-1 and SNU719 cells (Fig.2C). ChIPSeeker based annotation analysis identified 292 genes nearby 318 EBNA1 binding sites (Fig.2D). After that, we performed overlap analysis of EBNA1 ChIP-seq and VK1727 RNA-seq datasets to identify direct targets of VK1727. Ten in 27 differentially expressed genes were characterized as direct targets of VK1727, which were nearby EBNA1 binding sites (Fig.2D and Table S2). A conserved palindrome-like sequence (**GG**C**AG**CATATG**CT**G**CC**) was found in the center of EBNA1 binding sites (Fig.2D). CDC7 and POU2F1 were recognized as top two direct targets of VK1727 in PDX C15, and among the top targets in C666-1 and SNU719 (Fig.2D). EBNA1 binding sites were identified in CDC7 promoter and POU2F1 intron (Fig.2E and Table S2). CDC7 is known as a key regulator for initiation of DNA replication and promotion of G1/S or G2/M transition (40). CDC7 is centrally located in each of the regulatory networks perturbed by VK1727 (Fig.S1C, Fig. S2C and Fig.S3C). POU2F1 (also known as OCT1) is described as a stem cell transcription factor and functionalizes in self-renewal in context of colorectal cancer (41). Importantly, conservative EBNA1 binding sites were found in IL6R promoter (Fig.2E), which is consistent with a previous study from our lab (24).

**Figure 2.**
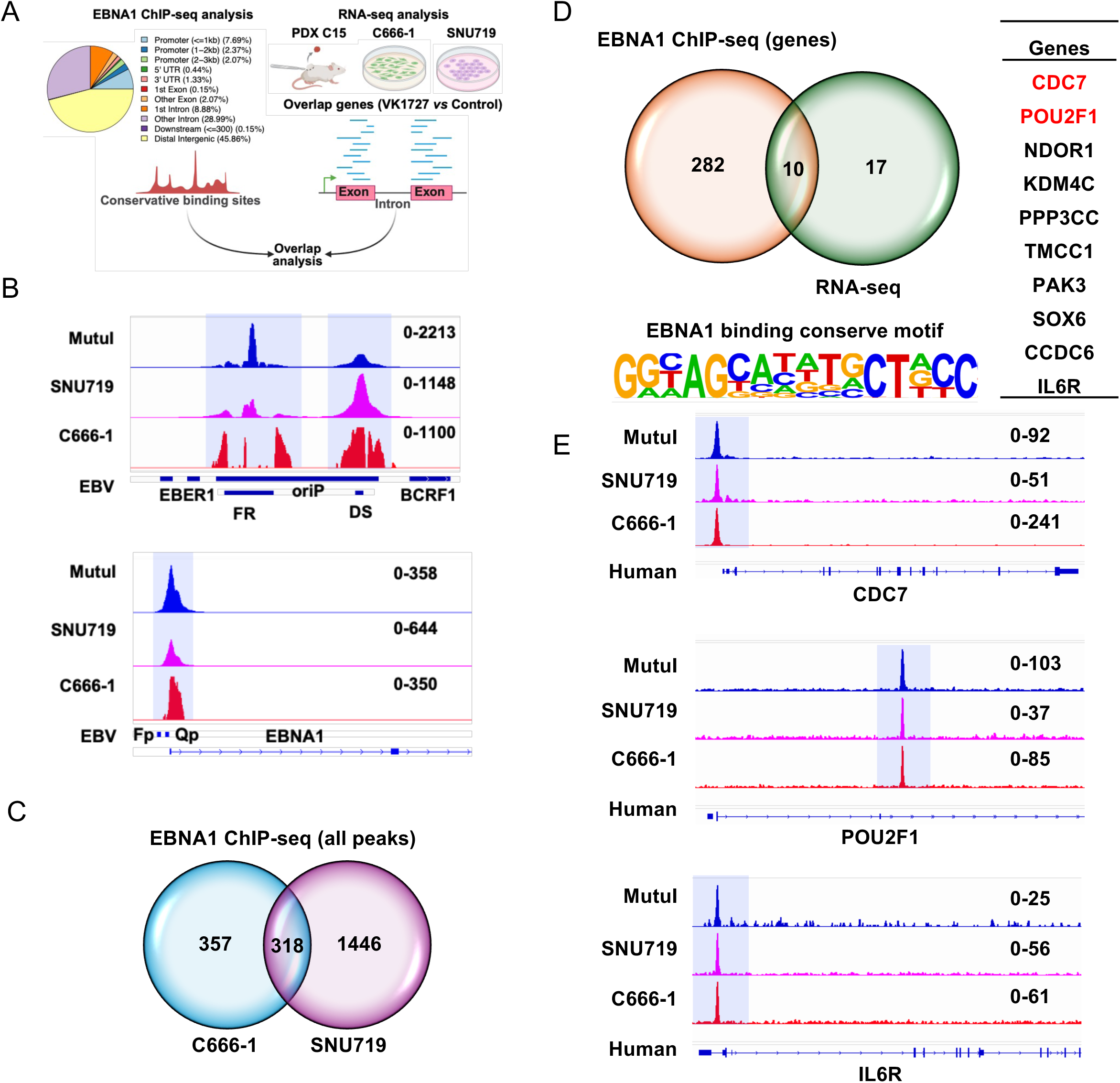
Overlap of EBNA1 ChIP-seq with VK1727 RNAseq. **A)** Schematic diagram for identification of VK1727 targets in differentially expressed gene sets among three tumor models. Potential targets of VK1727 were identified using conservative EBNA1 binding sites in viral and cellular genomes in SNU719 and C666-1 cells. Then overlap analysis was performed to identify differentially expressed genes nearby EBNA1 binding sites, which was achieved using *match* function in R program. **B)** EBNA1 ChIP-seq analysis of viral genomes in Mutu1, SNU719 and C666-1 cells. Conservative EBNA1 binding sites were characterized in FR, DS and Qp regions in viral genomes in Mutu1, C666-1 and SNU719 cells. **C)** EBNA1 conservative binding sites in cellular genomes between C666-1 and SNU719 cell lines. The bedtools were used to identify EBNA1 conservative binding sites between these two cell lines. 675 binding sites were identified in cellular genomes in C666-1 cells, while 1764 binding sites were characterized in cellular genomes in SNU719 cells. 318 conservative binding sites were enriched from overlap analysis. **D)** Overlap of EBNA1 ChIP-seq and RNA-seq in EBV^+^ epithelial cancer cells. Annotation of 318 conservative binding sites was achieved using ChIPseeker package in R program, which were nearby 292 cellular genes. Overlap analysis of EBNA1 ChIP-seq and RNA-seq characterized 10 direct targets of VK1727 in 27 differentially expressed genes. Motif analysis with HOMER identified a conservative palindromic-like sequence (**GG**C**AG**CATATG**CT**G**CC**) in the center of EBNA1 binding sites nearby 10 targets. **E)** EBNA1 binding sites in CDC7 promoter and POU2F1 intron. Conservative EBNA1 binding site was identified in prompter of CDC7 and intron of POU2F1 in Mutu1, C666-1 and SNU719 cells. Reliability analysis was confirmed using EBNA1 conservative binding site in promoter of IL6R.

In addition to the identification of CDC7 and POU2F1 as EBNA1 targets conserved in each of these cell and tumor types, we also characterized EBNA1 targets unique to one or more of these EBV epithelial cancer models. EBNA1 binding to CMKP2 and BMP4 was specific to SNU719 cells, while EBNA1 binding to MET was specific for C666-1 cells (Fig.S5, Tables S3, S4). These findings suggest that some EBNA1 engages both shared and cell type-specific genomic targets acrossEBV-associated epithelial cancers.

### ChIP-qPCR validation of VK1727 reduced EBNA1 binding to CDC7 and POU2F1 genes

We next conducted ChIP-qPCR assays to confirm VK1727 reduced EBNA1 binding to target regions. ChIP-qPCR assay showed VK1727 reduced over 70% of EBNA1 binding affinity to EBV FR region in PDX 15, C666-1 and SNU719 cells (Fig.3A-C), and more than 60% of EBNA1 binding at EBV DS and Qp regions in three tumor models (Fig.S6A-F). The oriLyt region served as a negative control showing only background levels of EBNA1 binding at this site identified in all three tumor models (Fig.S6G-I). These findings confirmed VK1727 efficiently reduced EBNA1 binding to known binding sites in the EBV genome. Next, we employed ChIP-qPCR to assay EBNA1 binding to the CDC7 promoter and POU2F1 intron in VK1727 treated PDX C15, C666-1 and SNU719 cells. We found that VK1727 reduced over 40% of EBNA1 binding affinity to POU2F1 intron (Fig.3D-F) and ∼80% of EBNA1 binding to CDC7 promoter in three tumor models (Fig.3G-I).

**Figure 3.**
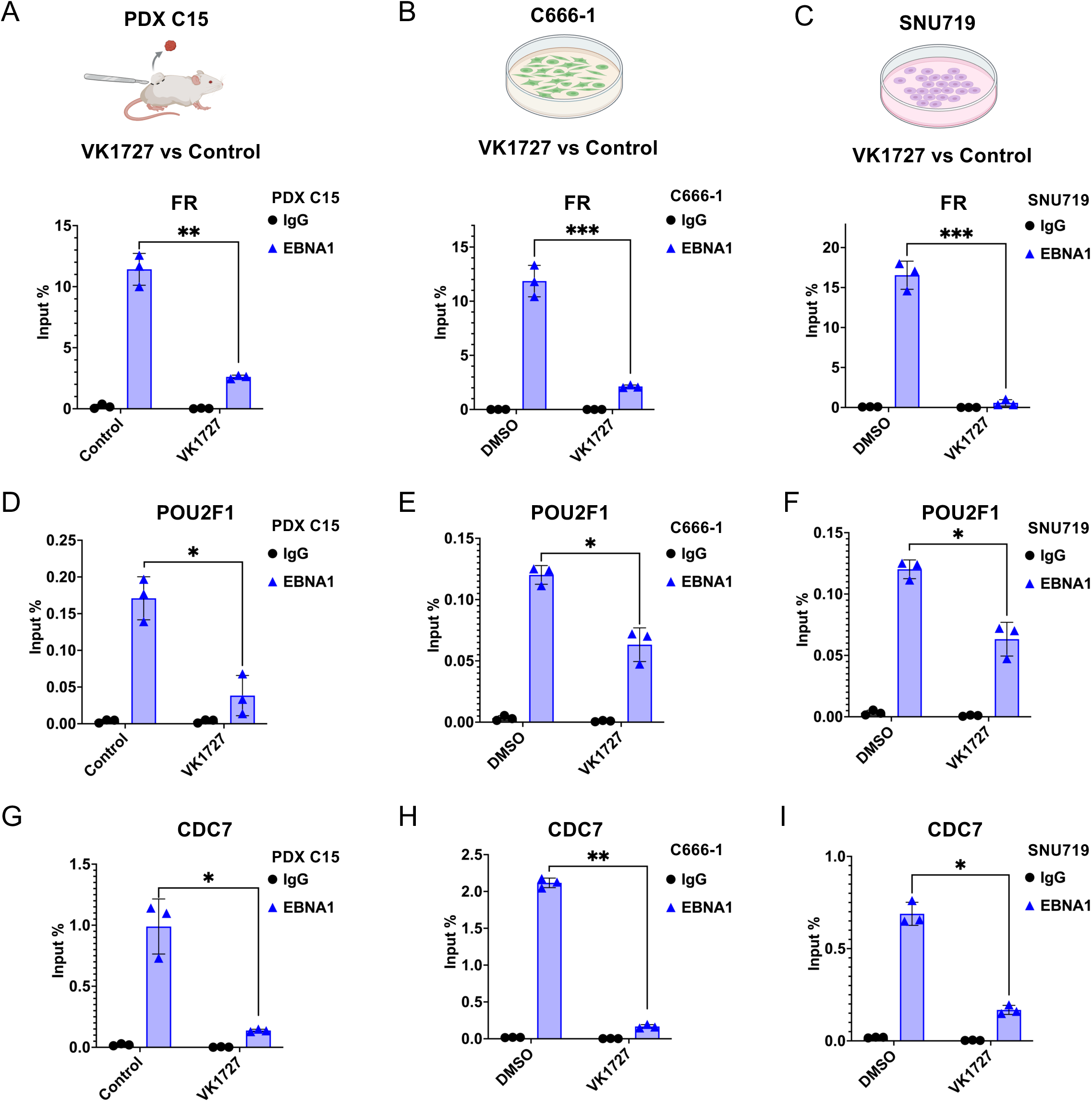
ChIP-qPCR validation of VK1727 reduced EBNA1 binding to CDC7 promoter and POU2F1 intron. **A-I)** ChIP-qPCR of EBNA1 or IgG binding to EBV FR region (A-C), POU2F1 intronic region (D-F), or CDC7 promoter (G-I) in PDX C15 (A, D, G), or C666-1 (B, E, H), or SNU719 (C, F, I). Error bars represent mean + SEM. Statistical comparisons between means were performed by Student’s t-test (2-tailed). *: p value <0.05, **: p value <0.01, ***: p value <0.005, ****: p value <0.001.

### RT-qPCR and Western blot validation of VK1727 reduction of CDC7 and Pou2F1 expression

Our results confirmed VK1727 reduces EBNA1 binding R CDC7 promoter and POU2F1 intron across all three tumor models. We next employed RT-qPCR assay and Western blot to detect expression of CDC7 and POU2F1 in VK1727-treated PDX C15, C666-1 and SNU719 cells (Fig. 4). RT-qPCR assay revealed a greater than 60% reduction of CDC7 transcription and ∼50% reduction of POU2F1 transcription after VK1727 treatments (Fig.4A-C). Western blot further confirmed a reduction of CDC7 and POU2F1 expression in VK1727 treated PDX C15, C666-1 and SNU719 cells (Fig.4D-F), which is consistent with transcriptomic analysis above. EBNA1 protein levels were not significantly altered at these relatively early timepoints, suggesting that VK1727 does not destabilize EBNA1 protein and that the observed effects on CDC7 and POU2F1 are likely direct, rather than an indirect consequence of loss of EBV episomes or altered expression of other viral genes. In addition, VK1727 treatments did not reduce transcription levels of CDC7 and POU2F1 in EBV– NPC HK-1 cells (Fig.S7), demonstrating that the effects of VK1727 treatment are specific to EBV+ cells.

**Figure 4.**
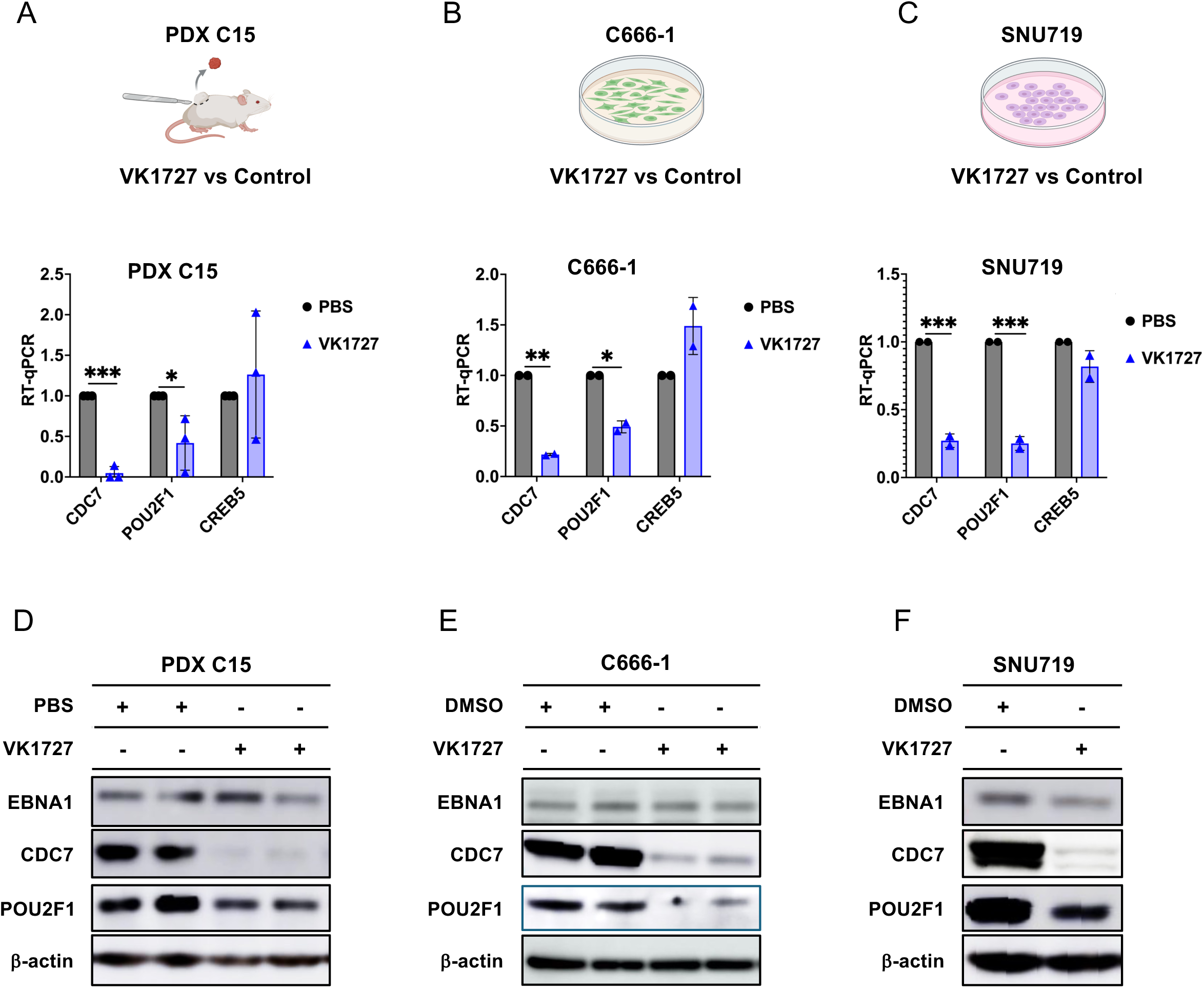
RT-qPCR and Western blot analysis of CDC7 and Pou2F1 after VK1727 treatment. **A-C)** RT-qPCR assay detected transcription of CDC7 and POU2F1 in PDX C15 after 5-day VK1727 treatments (A), C666-1 cells after 48-hour VK1727 treatments (B) and SNU719 cells after 48-hour VK1727 treatments (C). CREB5 was used as control. *GUSB* gene was used as internal control for normalization. **D-F)** Western blot probed for EBNA1, CDC7, POU2F1, and β-actin for PDX C15 (D) C666-1 cells (E) and SNU719 cells (F). β-actin was used as a loading control in western blot. Error bars represent mean + SEM. Statistical comparisons between means were performed by Student’s t-test (2-tailed). *: p value <0.05, **: p value <0.01, ***: p value <0.005, ****: p value <0.001.

### EBNA1 domains required for transcription regulation of CDC7 and Pou2F1

To gain insight into the mechanism by which EBNA1 regulates the transcriptional of CDC7 and POU2F1, we investigated the domain requirements of EBNA1. FLAG-tagged EBNA1 constructs, including full length lacking internal gly-ala repeats (WTΔGA), C-terminus deletion ΔC (lacking DBD) and C-terminus (DBD only), were overexpressed in EBV^-^ AGS cells, with an empty vector (EV) as a control. All EBNA1 proteins were expressed at comparable levels, as confirmed by FLAG-Western blot (Fig. 5B). Western blotting further demonstrated that CDC7 and POU2F1 protein levels were upregulated in AGS cells transfected with full-length WTΔGA EBNA1, but not in cells expressing either deletion mutant (Fig.5B). Consistent with this finding, RT-qPCR analysis showed significant upregulation of CDC7 and POU2F1 transcripts by full length EBNA1, but not by the deletion mutants (ΔC or C-terminus) (Fig.5C).

**Figure 5.**
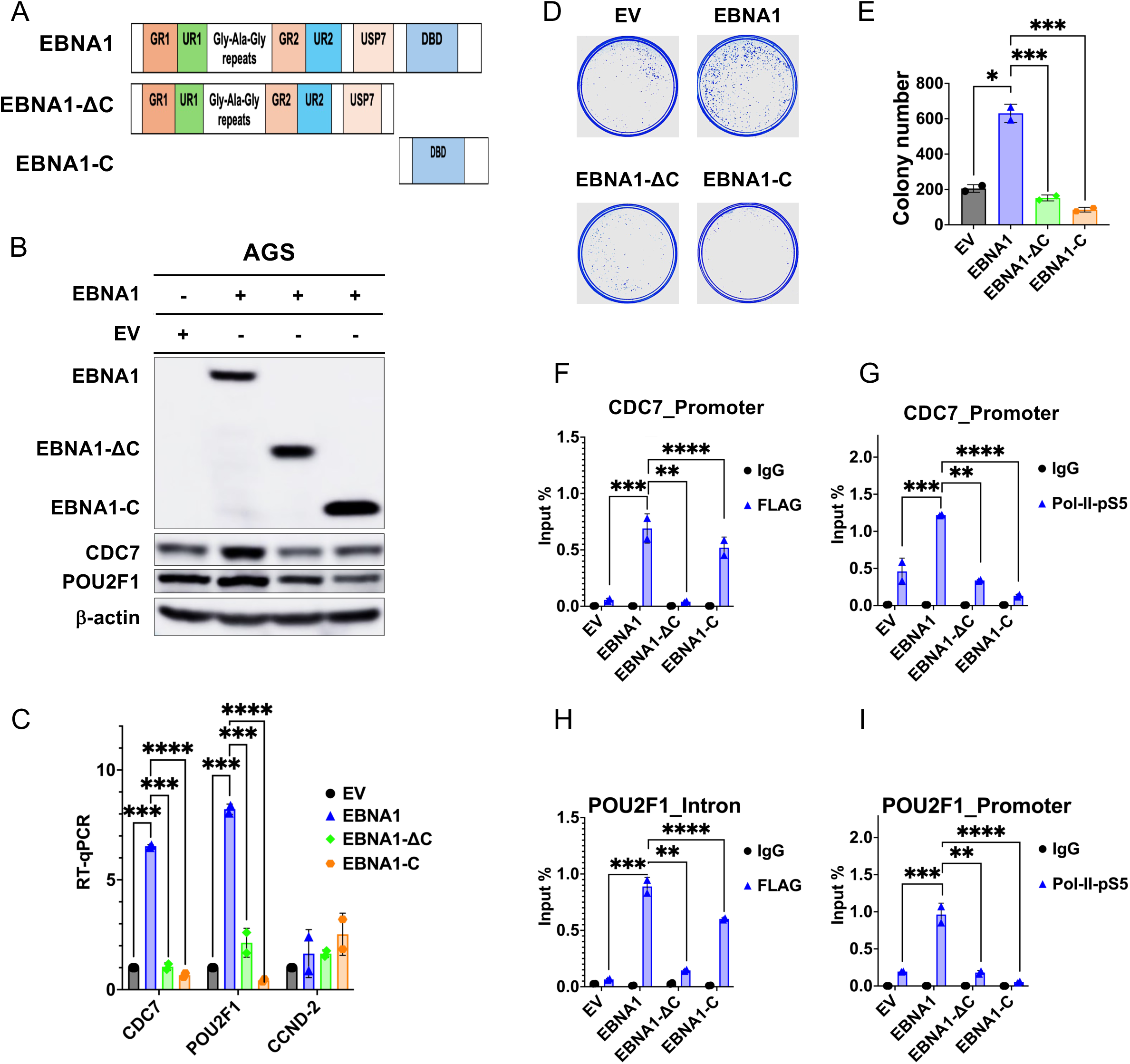
Structure-function analysis of EBNA1 regulation of CDC7 and Pou2F1. **A)** Schematic of EBNA1 domains and deletion mutants for FLAG-EBNA1, EBNA1-ΔC (aa1-459), and EBNA1-C (aa 460-641) for ectopic expression in AGS. **B)** Western blot analysis of AGS cells transfected with pCMV-FLAG empty vector (EV) or FLAG-EBNA1, EBNA1-ΔC, or EBNA1-C and probed for FLAG (top panel), CDC7, POU2F1, or β-Actin. Transfected AGS cells were collected for western blot analysis after 48-hour hygromycin B selection. **C**) RT-qPCR assay measuring RNA for CDC7, POU2F1, and CCND-2 in transduced AGS cells. *GUSB* gene was used as internal control for normalization. **D-E)** Colony formation assay detected proliferation of AGS cells transfected as in panel A and B. Quantification of colony forming assays shown in panel E. **F)** FLAG ChIP-qPCR assay for FLAG-EBNA1 binding to CDC7 promoter in transfected AGS cells. **G)** RNA PoI II**-**pS5 ChIP-qPCR assay at the CDC7 promoter in transfected AGS cells. **H)** FLAG ChIP-qPCR assay for FLAG-EBNA1 binding to POU2F1 intron in transfected AGS cells. **I)** RNA Pol II**-**pS5 ChIP-qPCR assay at POU2F1 promoter in transfected AGS cells. IgG control was used for each of the ChIP-qPCR assays. Error bars represent mean + SEM. Statistical comparisons between means were performed by Student’s t-test (2-tailed). *: p value <0.05, **: p value <0.01, ***: p value <0.005, ****: p value <0.001.

We next examined the functional consequences of EBNA1 on cellular proliferation using colony formation assays (Fig. 5D and E). Expression of full-length WTΔGA EBNA1 promoted colony expansion of AGS cells compared with empty vector, while overexpression of mutant EBNA1 reduced AGS cell proliferation relative to empty vector (Fig.5D and E). To further investigate the mechanism of transcription regulation, we performed FLAG and RNA Polymerase II phosphorylated at Ser5 (RNA Pol II-pS5) ChIP q-PCR analysis. RNA Pol II-pS5 is a critical marker of the transition from transcription initiation to elongation (42). FLAG ChIP-qPCR demonstrated WTΔGA and C-terminus of EBNA1 binding to CDC7 promoter and POU2F1 intron in transduced AGS cells, whereas EBNA1-ΔC showed no significant binding to these two target regions (Fig.5F and H). In contrast, RNA Pol II-pS5 ChIP q-PCR revealed that full length WTΔGA EBNA1 enhanced RNA PoII-pS5 recruitment to CDC7 and POU2F1 promoters in AGS cells compared with empty vector, while neither deletion mutant stimulated RNA Pol II-pS5 binding (Fig.5G and I). Together, these findings demonstrate that EBNA1 DNA-binding domain and N-terminal domains are both required for transcriptional activation of cellular target genes, and that recruitment of RNA Pol II-pS5 is a key component of this regulatory mechanism.

### Knockdown of POU2F1 impairs function of EBNA1 in EBV associated epithelial cancers

Our results demonstrated EBNA1 regulates the expression of POU2F1 and CDC7 through direct binding to their target regions. However, the functional role of POU2F1 in EBV+ epithelial cancers remains poorly understood. To address this, we employed small hairpin RNA (shRNA)–mediated knock down of POU2F1 in SNU719 cells. A pool of 3 shRNA achieved approximately 50% knock-down of POU2F1 expression (Fig. 6A). RT-qPCR analysis revealed that knockdown of POU2F1 resulted in significantly reduced transcription of EBV genes (EBNA1, LMP1, EBER1 and EBER2) and CDC7 in SNU719 cells (Fig.6A). Western blot analysis further confirmed that knockdown of POU2F1 resulted in a reduction of LMP1, EBNA1 and CDC7 protein levels in SNU719 cells (Fig.6B). In addition, colony formation assays demonstrated that knockdown of POU2F1 significantly impaired cellular proliferation and colony expansion in SNU719 cells (Fig. 6C and D). We next assayed whether POU2F1 influences EBV genome maintenance by measuring EBV DNA copy number using quantitative PCR. POU2F1 knockdown resulted in reduced copy number of EBV in SNU719 cells (Fig. 6E). These results show that knockdown of POU2F1 reduces EBV genome copy number and viral gene expression, including EBNA1, and supports a multifactorial role of POU2F1 in EBV+ epithelial cancers.

**Figure 6.**
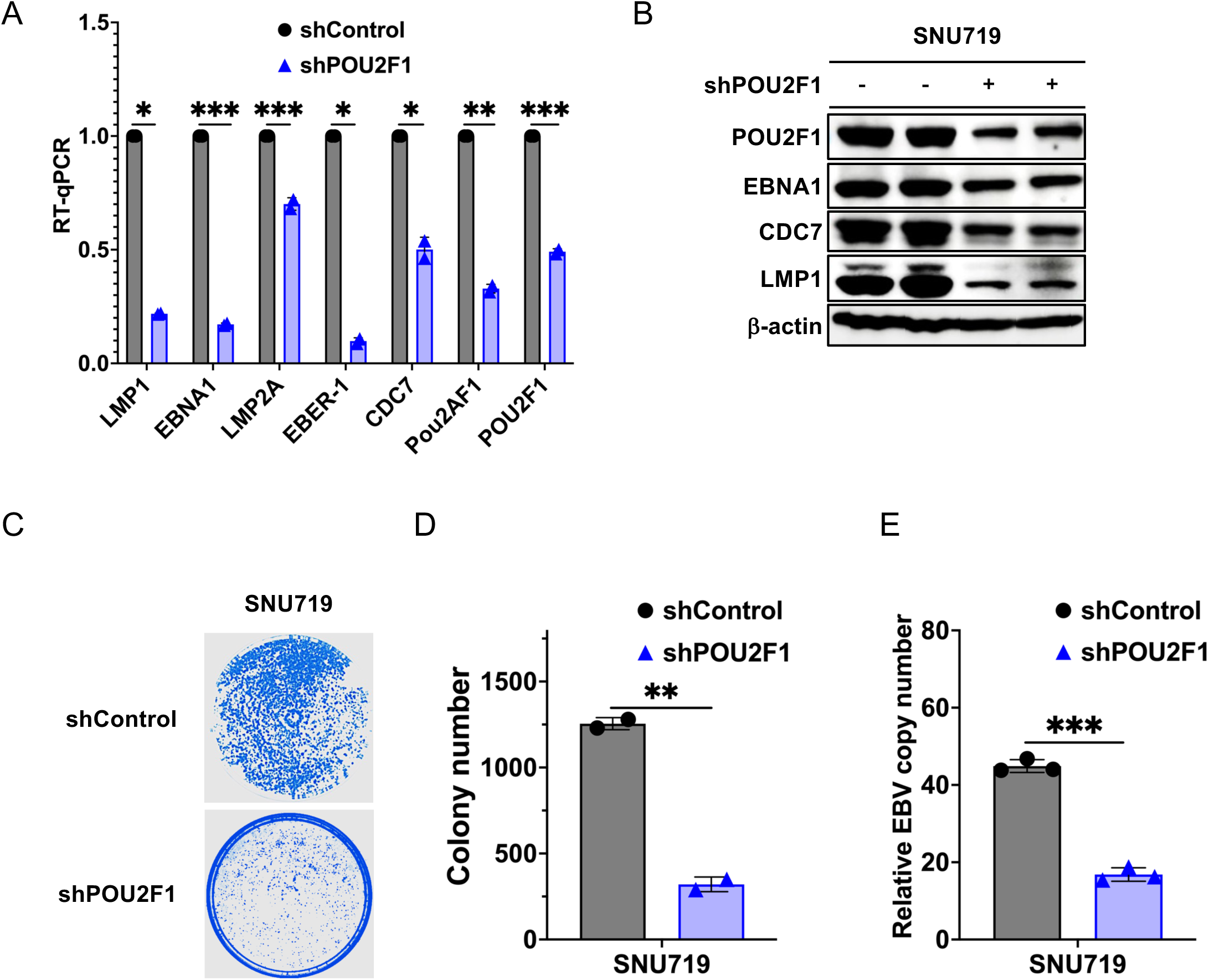
Pou2F1 knockdown reduces EBV gene expression and colony growth formation in SNU719 cells. **A)** RT-qPCR analysis of transcription of EBV genes (EBNA1, LMP2A, EBER-1, EBER-2, LMP1) and cellular genes for POU2F1, CDC7, Pou2AF in SNU719 cells after knockdown with shPOU2F1 or shControl. **B)** Western blot analysis of POU2F1, EBNA1, CDC7, LMP1 and β-Actin in SNU719 cells after knockdown with shPOU2F1 or shControl. **C-D)** Colony formation assay of SNU719 cells after knockdown with shPOU2F1 or shControl. Quantitation of colony formation shown in D. **E)** dd-qPCR assay measuring EBV DNA copy number in SNU719 cells after knockdown with shPOU2F1 or shControl. Error bars represent mean + SEM. Statistical comparisons between means were performed by Student’s t-test (2-tailed). *: p value <0.05, **: p value <0.01, ***: p value <0.005, ****: p value <0.001.

### CDC7 inhibitors phenocopy EBNA1 inhibitors to block cell proliferation in EBV+ epithelial cancer cell lines

To investigate the functional requirements of CDC7 in EBV+ epithelial cancers, we examined the selective effects of the CDC7 inhibitor Simurosertib on colony formation of EBV+ SNU719 cells compared to EBV– AGS epithelial cells (Fig. 7 and S8, S9). Colony formation assays shows that both of VK1727 and Simurosertib significantly inhibited colony expansion of SNU719 cells, while only modestly affecting colony growth in AGS cells (Fig.7A and B). Similar effects were observed in C666-1 (EBV+) and HK-1 (EBV–) cells (Fig.S9A and B). Western blot analysis revealed that Simurosertib markedly reduced expression of CDC7 in SNU719 cells but mildly affected CDC7 expression in AGS cells (Fig.7C). We also observed that VK1727 reduced both CDC7 and EBNA1 expression in SNU719 cells (Fig.7C).

**Figure 7.**
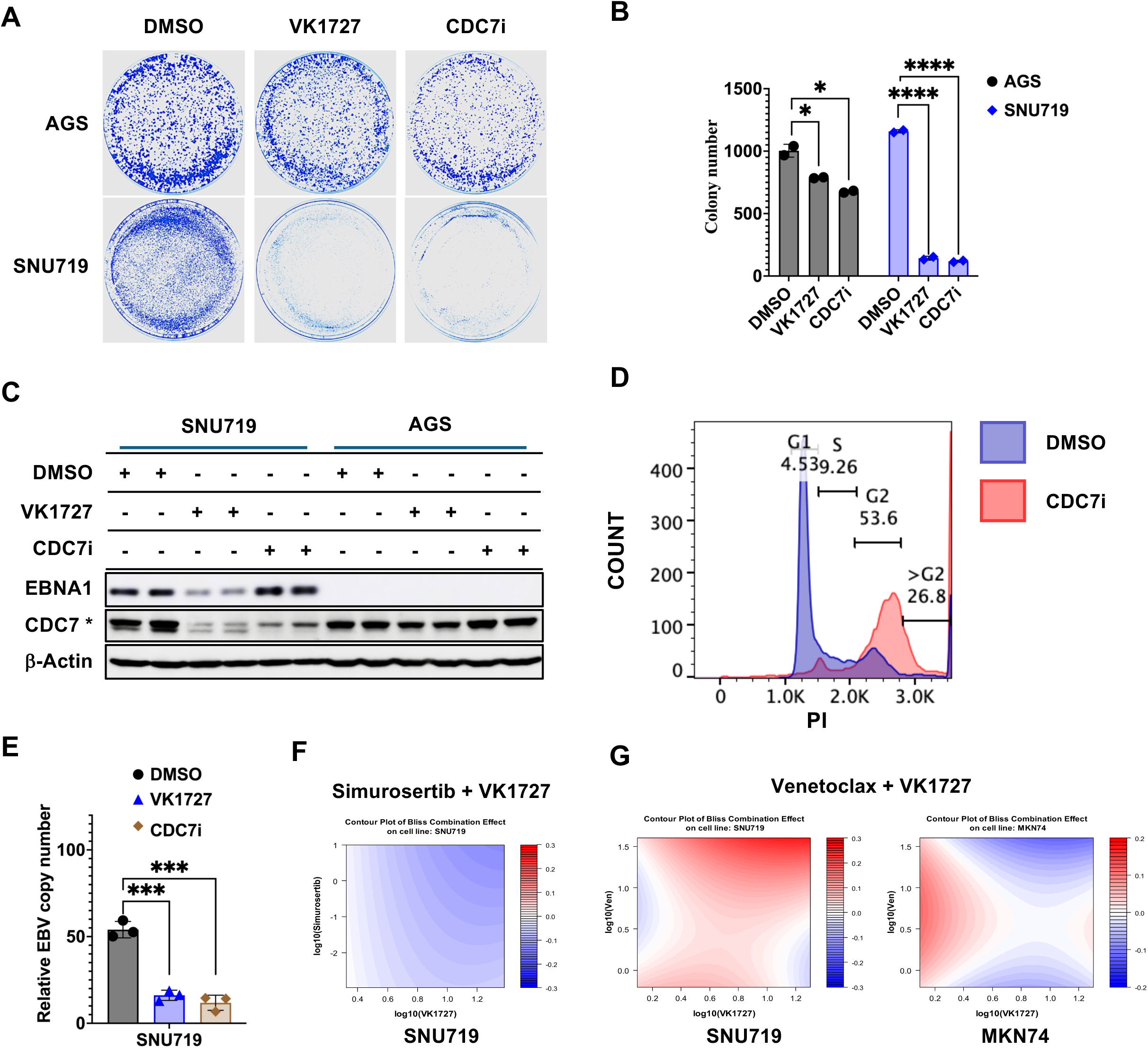
EBNA1 inhibitors are epistatic with CDC7 inhibitor Simurosertib and synergistic with Bcl2 inhibitor Venetoxlax. **A-B)** Colony formation of SNU719 cells after treatment with VK1727 (20 μM) or CDC7 inhibitor Simurosertib (1.5 μM). Error bars represent mean + SEM. Statistical comparisons between means were performed by Student’s t-test (2-tailed). *: p value <0.05, **: p value <0.01, ***: p value <0.005, ****: p value <0.001. **C)** Western blot analysis of EBNA1 and CDC7 in VK1727 or CDC7 inhibitor treated AGS and SNU719 cells. CDC7i treatment was 1.5 μM for 48 hours, VK1727 treatment was 20 μM for 48 hours. **D)** Flow cytometry analysis revealed G2/M arrest in SNU719 cells after CDC7 inhibitor treatments. SNU719 cells were treated with 1.5 μM CDC7 inhibitor at 48 hours. Propidium Iodide (PI) was employed to stain genomic DNA in SNU719 cells. **E)** dd-qPCR assay to measure copy number of EBV in SNU719 cells. Error bars represent mean + SEM. Statistical comparisons between means were performed by Student’s t-test (2-tailed). *: p value <0.05, **: p value <0.01, ***: p value <0.005, ****: p value <0.001. **F-G)** Synergy analysis of combination treatment with VK1727 and Simurosertib on SNU719 cell growth (F) or VK1727 and Venetoclax on SNU719 (left) and MKN74 (right) cell growth. Resazurin assay was used to measure cell viability under different combination treatment of VK1727 and Venetoclax. Bliss synergy score was calculated using SynergyFinder package (3.0) in R program.

Flow cytometry identified a G2/M cell cycle arrest in Simurosertib-treated SNU719 (Fig.7D). In some cells, Simurosertib appeared to impair chromosome segregation, resulting in an accumulation of cells with 2N DNA content (Fig.7D). We next measured EBV DNA copy number by DNA qPCR assay and observed a reduction of viral genome copy number in both VK1727 and Simurosertib treated SNU719 cells (Fig.7E). To determine whether Simurosertib and VK1727 act in the same pathway (epistatic), we assessed their combined effects using a two-stage response surface model. The combination of simurosertib and VK1727 did not exhibit synergistic growth inhibition in SNU719 cells (Fig.7F), which is consistent with both agents acting on overlapping or epistatic pathways.

In contrast, CDC7 inhibitors have been reported to act synergistically with BCL2 inhibitors in acute myeloid leukemia (AML) (43). We therefore tested whether VK1727 would similarly synergize with with the BCL2 inhibitor, venetoclax. VK1727 exhibited strong synergistic activity with Venetoclax EBV+ SNU719 cell lines (Fig. 7G, left), whereas no such synergy was observed in EB-negative MKN74 cells, suggesting that this effect is EBV-dependent (Fig. 7G, right). Together, these findings suggest that Simurosertib and VK1727 inhibit the same pathways controlling cell cycle progression and viral DNA replication in EBV+ epithelial cancers.

### EBNA1 regulation of CDC7 is conserved in EBV+ B-lymphoma

To determine if EBNA1 regulation of CDC7 in epithelial cancers is conserved in B-cell lymphoma, we analyzed RNA-seq from Mutu I cells treated with the EBNA1 inhibitor VK1850. VK1850 is a close analog of VK1727 and expected to function through an identical mechanism by binding EBNA1 and blocking its DNA binding. Comparative analysis across 4 treatment conditions identified three consistently upregulated genes (ARRDC3, DDIT3, NBPF1) (Fig.S10A) and five consistently downregulated genes (TK1, MCM7, CDK1, CDC7, RRM2) (Fig.S10B). RT-qPCR analysis confirmed that EBNA1 inhibition reduced transcription of CDC7 in MutuI cells following 72 hours of treatment (Fig.S10C). Pathway analysis did not identify common signaling pathways among the upregulated genes across treatments and cell types (Fig.S10D). However, cell cycle signaling pathway was identified among the downregulated gene sets across all four treatments and cell types (Fig.S10E). Together, these findings indicate that CDC7 and cell cycle pathways represent conserved targets of EBNA1 and are selectively downregulated upon EBNA1 inhibition in both epithelial and B-cell lymphoma contexts.

## DISCUSSION

EBV is a well-established human tumor virus due to its association with several human cancers, but its mechanisms of oncogenesis, particularly in epithelial cancers, are not fully understood (44). In EBV epithelial cancers, EBV episomes and EBNA1 protein are present in virtually all cancer cells (45, 46). EBNA1 is thought to make essential contributions to the oncogenic process by maintaining the viral episome and by binding cellular genomic sites to regulate host genes that are critical for oncogenic transformation (18, 47). In this study, we leveraged EBNA1 inhibitors as pharmacological tools to investigate the functional targets of EBNA1 in three different models of EBV-associated epithelial cancers. VK1727 and its closely related analog VK2019 target the DNA binding domain of EBNA1 (37, 39). Our findings further validate VK1727 as an inhibitor of EBNA1 DNA binding in both cell culture and PDX tumor models of nasopharyngeal carcinoma (e.g. PDX C15).

We analyzed the transcriptomic response to VK1727 in C666-1, SNU719, and PDX C15 and identified common genes and pathways that were differentially regulated by drug treatment. All three EBV+ epithelial cancer models shared cell cycle regulation, cellular senescence and transcriptional dysregulation in cancer as the top enriched pathways, with p53 signaling and EBV virus infection also scoring as highly significant. By integrating EBNA1 ChIP-seq data with known EBNA1 binding sites, we identified a small number of candidate genes with a high probability of being directly bound and transcriptional regulated by EBNA1. We further characterized the top 2 targets across all three epithelial cancer models, CDC7 and POU2F1, and demonstrated that EBNA1 binds and regulates these genes. Moreover, both genes make essential contributions to EBV+ epithelial cancer cell growth and survival.

Previous studies have identified EBNA1 bound transcriptional targets in lymphoid models of EBV cancer (24), including IL6R, MEF2B, and EBF1, each of which plays an important role in B-cell development and lymphomagenesis. In the present study, focusing on EBV+ epithelial cancer models identified IL6R as a significant dysregulated gene in SNU719, but not in nasopharyngeal carcinoma models. Neither MEF2D nor EBF1 scored as highly significant in epithelial cancers, suggesting that the action of EBNA1 is host cell or tumor type dependent.

Consistent with this idea, many EBNA1 regulated genes in NPC and EBVaGC were not shared between models and likely reflect cell type-specific regulation. EBNA1 DNA binding to host genes also varied across cell types, which may further contribute to differences in gene regulation. Many differentially regulated genes did not contain EBNA1 binding sites, likely reflecting indirect effects of EBNA1 inhibition on upstream transcriptional regulators. In addition, many EBNA1 binding sites were located in intergenic regions that could not be assigned to any specific gene, while some EBNA1-bound promoter sites did not correlate with transcriptional changes. Consequently, the overlap between EBNA1 binding and transcriptional deregulation, are relatively limited. While this may suggest that EBNA1 is not a strong transcriptional regulator compared to EBNA2 or EBNA3C, we nonetheless identified a distinct subset of EBNA1-bound genes that are differentially regulated. Here, we focused on the genes that are commonly regulated in the three epithelial cancer models to test the hypothesis that EBNA1 controls a set of essential, conserved targets in epithelial oncogenesis.

Our integrated data identified the cell cycle dependent kinase CDC7 and the stem cell transcription factor POU2F1 as direct targets of EBNA1-mediated gene regulation in all EBV epithelial cancer models examined (Fig.2E, Fig. 3G-I, Fig. 4, and Fig. 8). EBNA1 binding at the CDC7 promoter or the first intron of POU2F1 enhanced RNA Pol II-pS5 binding at the transcription start sites of these genes (Fig. 5), suggesting that EBNA1 contributes to the transcription activation. CDC7 has emerged as an attractive therapeutic target in several cancers, including pancreatic, prostate, breast, and lung cancers (48–51). CDC7 has a well-established function in replication origin firing by phosphorylating components of the MCM complex (Minichromosome Maintenance) required for the licensing and initiation of DNA replication (52, 53). MCM complex is also known to associate with EBV OriP and promotes viral replication (54, 55). Our study suggests CDC7 as a promising therapeutic target for EBV+ epithelial cancers, since CDC7 inhibitor simurosertib reduced colony expansion, and blocked G2/M progression in EBV+ epithelial cells (Fig. 7 and Fig. S9). Together with previous studies, our findings support a model that CDC7-mediated phosphorylation of the MCM complex contributes to the maintainance and persistence of EBV, consistent with the observation that simurosertib is epistatic with EBNA1 inhibition in reducing EBV copy number (Fig. 7E, Fig. 8).

**Figure 8.**
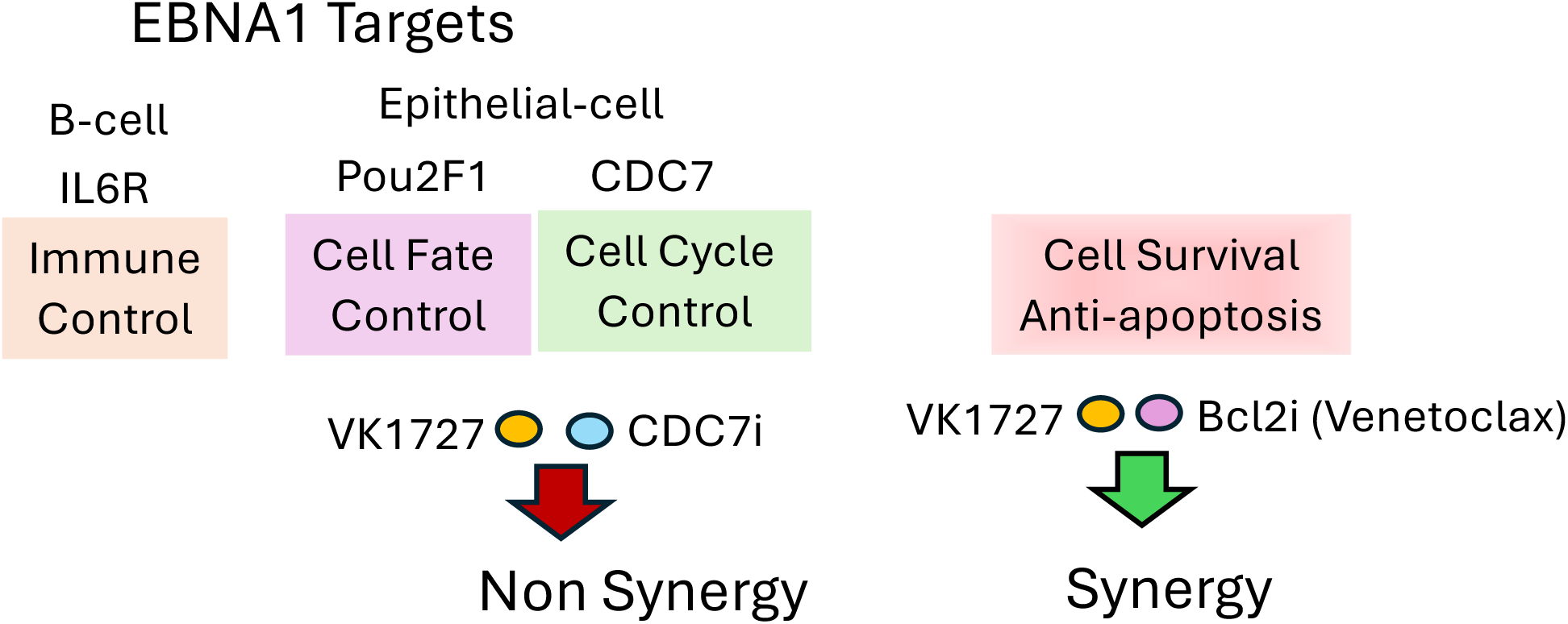
Model for EBNA1 directly bound cellular gene targets and drug synergy pathways. EBNA1 stimulates transcription of CDC7 to control cell cycle progression and activates expression of POU2F1 to regulate cell fate in EBV^+^ epithelial cancers. CDC7 inhibitors are non-synergistic and epistatic to EBNA1 inhibitors in epithelial cells. EBNA1 inhibitors are synergistic with Bcl2 inhibitors, which are not direct targets of EBNA1. Previous studies found EBNA1 regulated immune control genes in B-lymphocytes.

EBNA1 was also found to bind and regulate POU2F1 in each of the epithelial cancer models tested here. POU2F1 (also known as OCT1) has been reported to promote cancer stem self-renewal properties and resistance to chemotherapy (56). Our results suggest the POU2F1 pathway could be a therapeutic vulnerability in EBV+ tumors, as POU2F1 knockdown reduced cell proliferation, viral copy number and viral gene expression (Fig. 6). POU2F1 often cooperates with POU2F2 (known as OCT2) to stimulate transcription (57). OCT2 has been shown to bind to OriP in EBV genome and regulate transcription of EBV genes in EBV+ lymphoid cancers and transformed B-cells (58). These observations suggest that OCT1 and OCT2 may act cooperatively in B-cells, whereas OCT1, alone or in combination with OCT4, may be sufficient in EBV+ epithelial cancers to promote stem cell self-renewal and viral oncogenesis (59). Notably, OCT1 has also been investigated as target for cancer therapy and early-stage small molecule inhibitors have been developed for hepatocellular carcinomas (60).

EBNA1 is a multi-functional DNA binding protein that regulates viral replication through oriP and controls transcription of viral and host cellular genes through mechanisms that are not completely understood. Our study demonstrates that EBNA1-mediated transcriptional activation correlates with increases in RNA Pol II-pS5 occupancy at transcription start sites (Fig. 5).

Deletion of either the C-terminal DNA binding domain or the N-terminal tethering domains resulted in a loss of transcription activation function (Fig. 5). Previous studies have identified key functions of the EBNA1 N-terminal region in chromosome tethering (61), transcriptional activation (62), nuclear localization (63), as well as interactions with histone H1 (64, 65), BRD4 (66), USP7 (67), CK2 (68, 69), and RNAs (70, 71) including G-quadruplexes (72, 73). EBNA1 typically binds to 18-bp palindromic sequences in viral and cellular genome (74), though sequence variation may confer additional functional diversity. EBNA1 has been shown to induce replication fork stalling and double strand breaks at fragile chromosome sites enriched in EBNA1 binding sites (20). Although we did not observe transcriptional changes associated with this strong binding site in this study, it remains possible that distal genes may be affected. Future studies will be needed to map long-range EBNA1 interactions and better define their contribution to transcriptional regulation.

In conclusion, the studies presented here further demonstrate that EBNA1 inhibitors, such as VK1727, effectively block EBNA1 binding and perturb host gene expression through both direct and indirect mechanisms. Importantly, we have identified a conserved set of EBNA1 target genes across multiple EBV+ epithelial cancer models that may serve as targets for therapeutic intervention. Some of these EBNA1 targets may also be activated in EBV negative tumors through EBNA1-independent mechanisms. Our findings further suggest rational combination strategies, such as pairing EBNA1 inhibitors with Bcl2 inhibitors like venetoclax, which is also known to synergize with CDC7 inhibitors (43) (Figs. 7 and 8). EBNA1 inhibitors may also enhance the efficacy of CAR T-cell (75) and monoclonal antibody therapies (76) targeting additional cellular pathways identified in this study, offering new avenues for the treatment of EBV-associated epithelial cancers.

## MATERIALS AND METHODS

### Cell lines, patient derived xenograft C15 and EBNA1 inhibitor VK1727 treatments

Two EBV latently infected cell lines (C666-1, EBV+ nasopharyngeal carcinoma cell line; SNU719, EBV+ gastric cancer cell line) and two EBV- cell lines (HK-1, nasopharyngeal carcinoma cell line; AGS, gastric cancer cell line) were employed to explore versatile function of EBNA1 in EBV epithelial cancers. The four cell lines were cultured with Gibco™ RPMI 1640 Medium (Catalog number 11875101) at 37 ^°^C under 5% CO_2_ humidified incubators, which includes 10% fetal bovine serum (FBS; Gibco, Gaithersburg, MD, USA) penicillin and streptomycin (50 U/mL), and L-glutamine. In addition, patient derived xenograft (PDX) C15 was used as an *in vivo* model for elucidating critical role of EBNA1 in EBV+ epithelial tumor. PDX C15 was transplanted into NOD mice (NOD.*Cg-Prkdc^scid^ Il2rgtm1Wjl/ScJ*) for su(36)staining tumor growth *in vivo* according to previous description (36).

PDX C15 was generated as described previously (37). NOD mice (n=3) with engrafted PDX C15 tumors were treated with either vehicle control or 10 µM VK1727 injected every 12 hours in total of 5 days. C666-1 cells were treated with 10 µM VK1727 or DMSO control every 12 hours for a total of 2 days. SNU719 was treated with 20 µM VK1727 or DMSO control every 12 hours for a total of 2 days, as SNU719 cells maintain higher copy number of EBV episomes than C666-1 cells. Triplicates were performed for each condition (VK1727 and DMSO).

Cells were treated with CDC7 inhibitor Simurosertib (TAK-931; catalogue number, HY-100888) dose range 0.1-10 μM or Bcl2 inhibitor Venetoclax dose range of 0.6 to 40 μM to assess synergy with VK1727.

### RNA-seq, differentiation gene expression and functional analysis

Total RNAs were extracted from PDX C15, SNU719 and C666-1 cells using QIAGEN RNeasy Mini Kit (catalogue number, 74104), which were treated with VK1727 and DMSO. 1 µg total RNA of each sample was used for library construction using Illumina Stranded Total RNA Prep (Ligation with Ribo-Zero Plus) Kit (catalogue number, 20040529). Ribosome RNA depletion, RNA fragmentation, 3’ and 5’ end repairing, reversal transcription into cDNA, and adaptor ligation were achieved according to provided protocols (support.illumina.com). Then optimized PCR program (optimization of PCR cycles) was applied for amplification of cDNA libraries. The cDNA libraries were purified using AMPure XP Beads for DNA Cleanup (catalogue number, A63880) according to manufacturer protocols (support.illumina.com).

Quality of cDNA library was measured using 4200 TapeStation System (Agilent). High Sensitivity D1000 DNA ScreenTape assay (Catalog number, NC1786959) was used to quantify average size and concentration of cDNA library. Pair-end sequencing was conducted to identify sequences of cDNA reads in library by Genome center in Wistar Institute.

The fastaq files of RNA-seq raw datasets were obtained from FTP database provided by Genome center in Wistar Institute. Quality of RNA-seq raw datasets was determined using FastQC (version, 1.0.0) coupled with MultiQC. Adaptor sequences from each read was removed using Trim Galore (version 0.6.10) program with pair-end trimming algorithm. RNA reads were mapped to integrated human (Hg19, also known as GRCh37) and EBV (NC007605) reference genomes using STAR aligner (version 2.7.11b). The count table of reads per gene was generated using featureCounts program (77) and utilized for subsequential analysis (*e.g.* Differential Gene expression and functional analysis). DESeq2 package (version, 1.50.0), embodied in R (version, 4.5.2) program, was used for differential gene expression analysis (78). The EnhancedVolcano and heatmap.2 packages were employed to generate volcano plots and heatmaps for visualization (79). The clusterProfiler package was applied to decode biological function of differentially expressed genes in KEGG (Kyoto Encyclopedia of Genes and Genomes) database (80). The DOSE package was utilized to generate bar plot and network of enriched biological function (81). The *match* function in R program was used to identify overlapping targets of downregulated and upregulated genes among PDX C15, C666-1 and SNU719 under VK1727 treatments.

In addition, we performed comparative transcriptomic analysis of RNA-seq datasets across four EBNA1 inhibitor treatments (VK1727: SNU719, C666-1 and PDX C15) and Mutu1 (VK1850: Burkitt lymphoma cell line) cells. Differential expression analysis was conducted using DESeq2 on raw count data, where genes with fewer than 10 counts were filtered out and significance was defined by an adjusted p-value < 0.05 (78). To identify shared and unique gene sets, intersections were visualized using 4-way Venn diagrams via the VennDiagram package (82). Global functional profiling was performed across all groups using the compareCluster function in clusterProfiler for KEGG pathway enrichment (83).

### Overlapping analysis between EBNA1 ChIP-seq and RNA-seq datasets

EBNA1 ChIP-seq fastaq files of SNU719 and MutuI cells were downloaded from FTP database provided by Genome Center in Wistar Institute, which was already published from our lab (GSE289710) (30). EBNA1 ChIP-Seq of C666-1 cells was fulfilled in consistency with previous description (30). The fastq files of EBNA1 ChIP-seq raw datasets were retrieved from FTP database provided by Genome Center in Wistar Institute. Quality control and trimming steps were conducted using FastQC and Trim Galore (version 0.6.10) programs. BowTie2 was employed to map DNA reads to the integrated human (Hg19) and EBV (NC007605) reference genomes using reported protocols (24). Samtools and Picard were used to remove duplicate reads in BAM files. MACS2 program was utilized to call EBNA1 peaks in cellular (Hg19) and viral (NC007605) genomes (24). ChIPSeeker package, embodied in R program, was used to annotate peaks in narrow-peak files to cellular genomes and generate peak annotation table (84). Then overlap analysis were performed to match annotated peaks with commonly upregulated or downregulated genes under VK1727 treatments, which employs *match* function in R program.

### ChIP assay for identifying EBNA1 and RNA polymerase II CTD phosphorylated Ser5 binding to viral and cellular genomes

10^6^ C666-1 or SNU719 cells were seeded on six-well plates. Three wells of C666-1 or SNU719 cells were treated with 0.3 % DMSO, the other three was dealt with 10 or 20 µM VK1727 every 12 hours in total of 2 days. Treated C666-1 or SNU719 cells were collected for ChIP assay using Gibco™ RPMI 1640 Medium (Catalog number 11875101). ChIP assay was accomplished based on reported protocols from our lab (30). Q-PCR was used to decode EBNA1 binding to FR, DS, Qp and OriPLyt in viral genome using identified primers (30).

Primers were designed to verify EBNA1 binding to promoter region of CDC7 and first intron region of POU2F1 in EBV epithelial cancers (ChIP-EBNA1forward primer for CDC7, AACCCACCTACCTCATAGCC; ChIP-EBNA1reverse primer for CDC7, TTCCTTTTCGTTGAGTGCCC; ChIP-EBNA1 forward primer for POU2F1, GCCAAGCTCACTCACACAG; ChIP-EBNA1 reverse primer, TCTGGCTCTTCTCATGTGCT).

Q-PCR running program were conducted in accordance with previous description (30). Input percentage was utilized to determine differences of EBNA1 binding affinity to viral and cellular genomes between VK1727 treatment and control groups. Input percentage was calculated using 2 ^-ΔCT^. The ΔCT was generated through differences of CT value between IP samples (IgG or EBNA1) and Input samples. Moreover, ChIP assay was employed to identify EBNA1 binding to genomic regions of interests in control and VK1727 treated C15 tumors.

ChIP-qPCR followed previously described protocols (85). Primers were designed to target regions close to TSS (from -1000 bp to 1000 bp) of CDC7 and POU2F1 (ChIP Pol II-pS5 forward primer for CDC7, GTTTCCGACGGTTTGTTCCA; ChIP Pol II-pS5 reverse primer for CDC7, AGAGACCGAACCAGATGCTT; ChIP Pol II-pS5 forward primer for POU2F1, CAGAGCGAGGGAGGGTTTAT; ChIP Pol II-pS5 reverse primer, AGCCGGGGTTGAGTATGAAT). ChIP assay was conducted using antibody against to RNA Pol-II pS5 (Catalogue number, 2687451; Active Motif). Input percentage was utilized to determine differences of RNA Pol- II PS5 access to promoter of CDC7 and POU2F1 between control and different treatments.

### Transient transfection of wild type, C-terminus deletion ΔC, and C-terminus of EBNA1 in AGS cells

Wild type full length lacking internal gly-ala repeats (WTΔGA), C-terminus deletion ΔC (lacking DBD, deletion from 460 to C-terminus 641) and C-terminus (DBD only, deletion from N terminus 2 to 459) of FLAG-EBNA1 with OriP have been described previously (86). The empty vector pCMV-FLAG (pPL748) were used as control. Hygromycin B resistant gene was cloned into backbone of four plasmids. Four plasmids were transiently transduced into AGS cells using Lipofectamine 2000 kit (Catalog number 11668027, Invitrogen). 200 µg/mL hygromycin B (Catalog number 10687010, Gibco™) was used to select AGS cells with overexpression of wild type, ΔC, and C-terminus of FLAG-EBNA1.

### POU2F1 knockdown and selective effects of CDC7 inhibitors on EBV+ epithelial cancers

Short hairpin RNAs (shRNAs) were employed to knock down POU2F1 in SNU719 cells. The shRNAs were purchased from the Wistar Institute collection of Sigma shRNA library (Full sequences for shRNA-1, CCGGGCAAAGGAGAGAAGGGAGAAACTCGAGTTTCTCCCTTCTCTCCTTTGCTT TTT; shRNA-2, CCGGCCAAACTACCATCTCTCGATTCTCGAGAATCGAGAGATGGTAGTTTG GTTTTT; shRNA-3, CCGGGCTGTGACGAATCTTTCAGTTCTCGAGAACTGAAAGATTCGTCACAGCTTTTT). Lentivirus production and purification were performed as described previously (87). Purified lentivirus, cloned with POU2F1 shRNAs or scramble control, was recruited to infect SNU719 cells for 48 hours, which was cultured with Gibco™ RPMI 1640 Medium (supplemented with 10 µg/mL polybrene, catalogue number K2701). 2 µg/mL puromycin (Catalog number A1113803, Gibco™) selection was utilized to obtain POU2F1 knockdown pool of SNU719 cells. RT-Q-PCR assay and western blot were conducted to quantify expression of POU2F1 and other genes of interests in control and POU2F1 knockdown SNU719 cells.

### Resazurin assay, Colony formation and Synergy analysis

Resazurin viability assay was performed as described previously (86). Briefly, 8000 EBV+ and EBV- cancer cells were seeded into each well in 96-well plate. Different concentrations of Simurosertib (100 nM-10 μM) were used to treat with EBV+ and EBV- cancer cells on 9 days. Resazurin sodium salt (catalogue number, R7017-1G) was used to identify cell viability. Optical value of cell viability was obtained using PerkinElmer EnVision Xcite multilabel plate reader. Puromycin Dihydrochloride (Catalog number A1113803) was utilized as positive control to normalize inhibition of cell viability. The percentage of growth inhibition was calculated in accordance with previous study (37).

For colony formation assays, 2000-3000 cells of EBV^+^ and EBV^-^ epithelial cancer cells were seeded on 60 mm × 15 mm Corning® tissue-culture treated culture dishes (CLS430166-500EA) for 7 days. Then 10 µM VK1727 and 1.5 µM (Selective dose) Simurosertib were prepared to treat cell clones on 50 × 15 mm Petri Dishes every 12 hours in total of 7 days. 0.3 % DMSO was used for control group. Duplicates were conducted for each condition (VK1727 and Simurosertib treatments, or POU2F1 knockdown). After that, cell clones were fixed with 10 % formaldehyde (SKU: C3966-1L) at 10 mins. The fixed cell clones were washed with PBS and stained with 0.1 % crystal violet (Catalogue number, V5265-2; Sigma) at 10 mins. Biorad ChemiDoc MP Imaging System was used to scan cell clones on 50 × 15 mm Petri Dishes and generate TIFF files for quantitative analysis. Image J program (version 1.54g) was used to determine number of cell clones on 50 × 15 mm Petri Dishes. Student’s T test was used to identify numeric difference of cell clones between control and treatments using Prism-GraphPad (Version 10). P value was utilized to measure significance of difference between control and treatments.

Synergy analysis of VK1727 and other inhibitors (Simurosertib and Bcl2 inhibitor Venetoclax) were performed using resazurin viability assay. Combinations of different concentrations of VK1727 (0.1-20 μM) with simurosertib (0.1-10 μM) or venetoclax (0.6 to 40 μM) were employed to inhibit cell viability of SNU719 (EBV^+^) and MKN74 (EBV^-^) cells. The percentage of growth inhibition was calculated in accordance with previous study (37). SynergyFinder package (3.0) in R program was utilized to analyze synergy scores between two inhibitors using two-stage response surface model (88).

### Western blot

Total protein was extracted from cells using Radioimmunoprecipitation assay buffer (RIPA, Thermo Fisher Scientific) supplemented with proteinase inhibitor (Roche cOmplete^TM^ Mini Proteinase Inhibitor Cocktail Tablets). Goat anti mouse or rabit IgG conjugated with Horseradish Peroxidase (HRP) (Catalogue number, 5178-2504 or 644005; Bio-Rad) was used to amplify intensities of blots. Immobilon Crescendo Western HRP substrate (WBLUR0500, Millipore) and Biorad ChemiDoc MP Imaging System were employed for detection of immunoblot on 0.2 µm nitrocellulose membrane (Catalogue number, 1620112; Bio-Rad). Western blots were probed with affinity purified Rabbit antibody against to EBNA1 generated in-house, or commercial antibodes to POU2F1 (Catalogue number, ab272867; Abcam), CDC7 (Catalogue number, sc-56275; Santa Cruz Biotechnology) .

### RT-qPCR Assay

Total RNAs were extracted from PDX C15, SNU719 and C666-1 cells using QIAGEN RNeasy Mini Kit (catalogue number, 74104), which were treated with VK1727 or DMSO. 2 μg total RNA was used to synthesize cDNA using SuperScript™ IV First-Strand Synthesis System (Catalogue number, 18091300). The cDNA products were diluted by 10 times for quantitative PCR analysis. Power SYBR™ Green PCR Master Mix (Catalog number 4368708) was used for quantitative PCR assay. The qPCR was performed with provided program (95 ℃ for 30 s; 45 cycles: 95 ℃ for 30 s, 60 ℃ for 10 s, 72 ℃ for 10 s; 65 ℃ to 95 ℃ at 0.1 ℃/s for melting curve analysis) in QuantStudio 6 and 7 Pro Real-Time PCR Systems. *GUSB* was utilized as a reference gene for normalization. Relative fold change value was calculated by 2^−ΔΔ𝐶𝑡^ Method. Δ𝐶𝑡 = Ct (target gene)- Ct (*GUSB*), and ΔΔ𝐶𝑡 = Δ𝐶𝑡 (Control) - Δ𝐶𝑡 (treatment).

### Flow cytometry for detection of cell cycle

Flow cytometry for measuring cell cycle of EBV^+^ and EBV^-^ epithelial cancer cells used Propidium Iodide (PI) based cell cycle detection as previously described (37). FlowJo™ (version) was utilized to identify G1-S or G2-M transition events of EBV^+^ and EBV^-^ epithelial cancer cells under VK1727 and Simurosertib treatments. Then Student’s T test was used to identify numeric difference of these events between control and treatments using Prism-GraphPad (Version 10). P value was employed to measure significance of difference between control and treatments.

### Relative quantitative PCR (qPCR) assay for identification of EBV copy number

Relative DNA qPCR (ddPCR) was utilized to determine copy number of EBV episomes in EBV^+^ epithelial cancer cells (SNU719 and C666-1) using previously described methods (30). Briefly, genome DNAs were extracted from treated 1x10^6^ SNU719 and C666-1 cells using QIAamp DNA Blood Mini Kit (Catalogue number, 51104; Qiagen). 30 ng DNA from each sample was sonicated into short size (200 bp to 600 bp) using Bioruptor® Plus sonication device (catalogue number, B01020014). Then sonicated DNA was utilized for detection of EBV copy number. EBV oriLyt (30) and GAPDH promoter regions (89) were employed to determine copy number of EBV episomes using identified primers. Relative EBV copy number was calculated using 2 ^- ΔCT^. The ΔCT was generated through differences of CT value between EBV OriLyt and GAPDH promoter. Then Student’s T test was used to identify difference of EBV copy number between control and treatments using Prism-GraphPad (Version 10). P value was employed to measure difference significance between control and treatments.

### Data accessibility

RNA-seq raw datasets (FASTQ files) were deposited in Gene Expression Omnibus database in NCBI. The accession number for deposited datasets were GSE315867 (RNA-seq datasets) and GSE315844 (ChIP-seq datasets).

## Acknowledgements

We thank members of the Wistar Cancer Center Cores in Genomics and Bioinformatics for their excellent technical support. This work was supported by grants from NIH P01 CA269043, R01 CA093606, R01 DE017336 to PML and CA259171 (TEM) and P30 Cancer Center Support Grant P30 CA010815 to the Wistar Institute (D. Altieri).

